# Association of HLA class I type with prevalence and outcome of patients with acute myeloid leukemia and mutated nucleophosmin

**DOI:** 10.1101/411645

**Authors:** Kateřina Kuželová, Barbora Brodská, Johannes Schetelig, Christoph Röllig, Zdeněk Ráčil, Juliane Stickel Walz, Grzegorz Helbig, Ota Fuchs, Milena Vraná, Pavla Pecherková, Cyril Šálek, Jiří Mayer

## Abstract

Acute myeloid leukemia with mutated nucleophosmin (NPMc+ AML) forms a distinct AML subgroup with better prognosis which can potentially be associated with immune response against the mutated nucleophosmin (NPM). As the T-cell-mediated immunity involves antigen presentation on HLA class I molecules, we hypothesized that individuals with suitable HLA type could be less prone to develop NPMc+ AML. We compared HLA class I distribution in NPMc+ AML patient cohort (398 patients from 5 centers) with the HLA allele frequencies of the caucasian population and found HLA-A*02, B*07, B*40 and C*07 underrepresented in the NPMc+ AML group. Presence of B*07 or C*07:01 antigen was associated with better survival in patients without concomitant *FLT3* internal tandem duplication. Candidate NPM-derived immunopeptides were found for B*40 and B*07 using prediction software tools. Our findings suggest that a T-cell-mediated immune response could actually explain better prognosis of NPMc+ patients and provide a rationale for attempts to explore the importance of immunosuppressive mechanisms in this AML subgroup.

## Introduction

In relation with several breakthrough concepts, such as the introduction of immune checkpoint inhibitors or the construction of T-cells with chimeric antigen receptors, the role of the immune system in prevention and treatment of leukemias is currently intensively studied. The curative potential of hematopoietic cell transplantation indicates that the activity of the immune system is important also in acute myeloid leukemia (AML). However, the immune response is often inhibited by a number of different mechanisms and although they are explored primarily in solid malignancies, at least some of them are relevant also for AML [1-3]. Antibodies against inhibitory receptors, such as ipilimumab, nivolumab and pembrolizumab, are efficient in blocking selected immunosuppressive processes and are tested in clinical trials also for leukemias. *In vitro* studies show that the immune checkpoint inhibition significantly increases the potential of cytotoxic T-cells to lyse leukemia blasts [4,5]. One of the remaining obstacles in the development of protocols for adoptive immunotherapy is the absence of suitable antigens. In the case of AML, pan-leukemic markers such as WT1, PRAME or Aurora kinase are usually considered [6]. C-terminal mutations in nucleophosmin are characteristic for AML, they are found in about a third of AML patients and are relatively stable during the disease course [7]. Patients with these mutations form a specific group with better prognosis which might be associated with an immune response against the mutated NPM1 or against its interaction partners [8,9].

To test the hypothesis that T-cell mediated immune response contributes to disease control in patients with *NPM1* mutation, we compared the distribution of HLA class I allelic groups between patients with *NPM1*-mutated AML and the normal population and we searched for an impact of the HLA-repertoire of the patient on survival outcomes.

## Results

We have previously performed a single-center pilot analysis of HLA class I distribution in patients with acute myeloid leukemia (AML, N = 63). The study indicated that the frequency of several HLA allelic groups was lower in patients with nucleophosmin mutations (denoted as NPMc+ in association with the cytoplasmic localization of the mutated protein) compared to the normal values as well as to the frequency observed in AML patients with wild-type *NPM1* [10]. In the present study, we analyzed HLA distribution in a larger cohort (398 patients with *NPM1* mutation from 5 centers). The basic parameters of the merged cohort and of the relevant subcohorts are given in Table 1.

**Table 1.** Basic parameters of patient cohorts

|  | age at diagnosis (median, range) | Flt3-ITD (% of positive cases) | Sex (% of females) | mutA/D (%) | Relapsed (%) | Transplant (%) | Karyotype aberration |
| --- | --- | --- | --- | --- | --- | --- | --- |
| all (N = 398) | 51.0 (19 - 73) | 42.7 | 54 | 92.6 | 37.5 | 39.2 | 18 of 133 |
| A02+ (N = 164) | 51.5 (21 - 70) | 41.5 | 54 | 90.4 | 37.2 | 37.0 | 9 of 62 |
| B07+ (N = 81) | 53.0 (19 - 73) | 30.5 | 56 | 93.4 | 34.2 | 35.5 | 2 of 22 |
| B40+ (N = 25) | 50.5 | 48.0 | 60 | 87.5 | 37.5 | 37.5 | 3 of 9 |
|  | (30 - 68) |  |  |  |  |  |  |
| C07+ (N = 134) | 53.0<br>(19 - 73) | 45.2 | 53 | 94.5 | 43.4 | 40.9 | 9 of 53 |
| Prague<br>subcohort (N =<br>94) | 52.5<br>(24 - 68) | 45.5 | 60 | 92.6 | 45.7 | 54.8 | 15 of 77 |
| C*07:01+<br>Prague (N =<br>20) | 52.0<br>(24 - 67) | 50.0 | 65 | 95.0 | 46.7 | 53.6 | 4 of 18 |

Basic characteristics of NPMc+ AML patient cohort and of its subgroups involving patients with selected HLA class I alleles as indicated. The Prague subcohort and its C*07:01-positive subgroup are given in the last two rows. Nucleophosmin mutations were of type A/D (both A and D result in the same protein product) or of type B. Flt3-ITD, internal tandem duplication in FLT3 gene. Karyotype information from cytogenetic analysis was available only in a part of patients and is given as the number of cases with karyotype aberrations as a part of all cases with available results.

The incidence of the HLA allelic groups A’02, B’07, B’40 and C’07 were found to be significantly lower in AML NPMc+ patients compared to the normal frequency obtained from www.allelefrequencies.net for caucasian populations (Fig 1). On the other hand, the frequency of A’24 was higher in the AML NPMc+ group.

**Figure 1:**
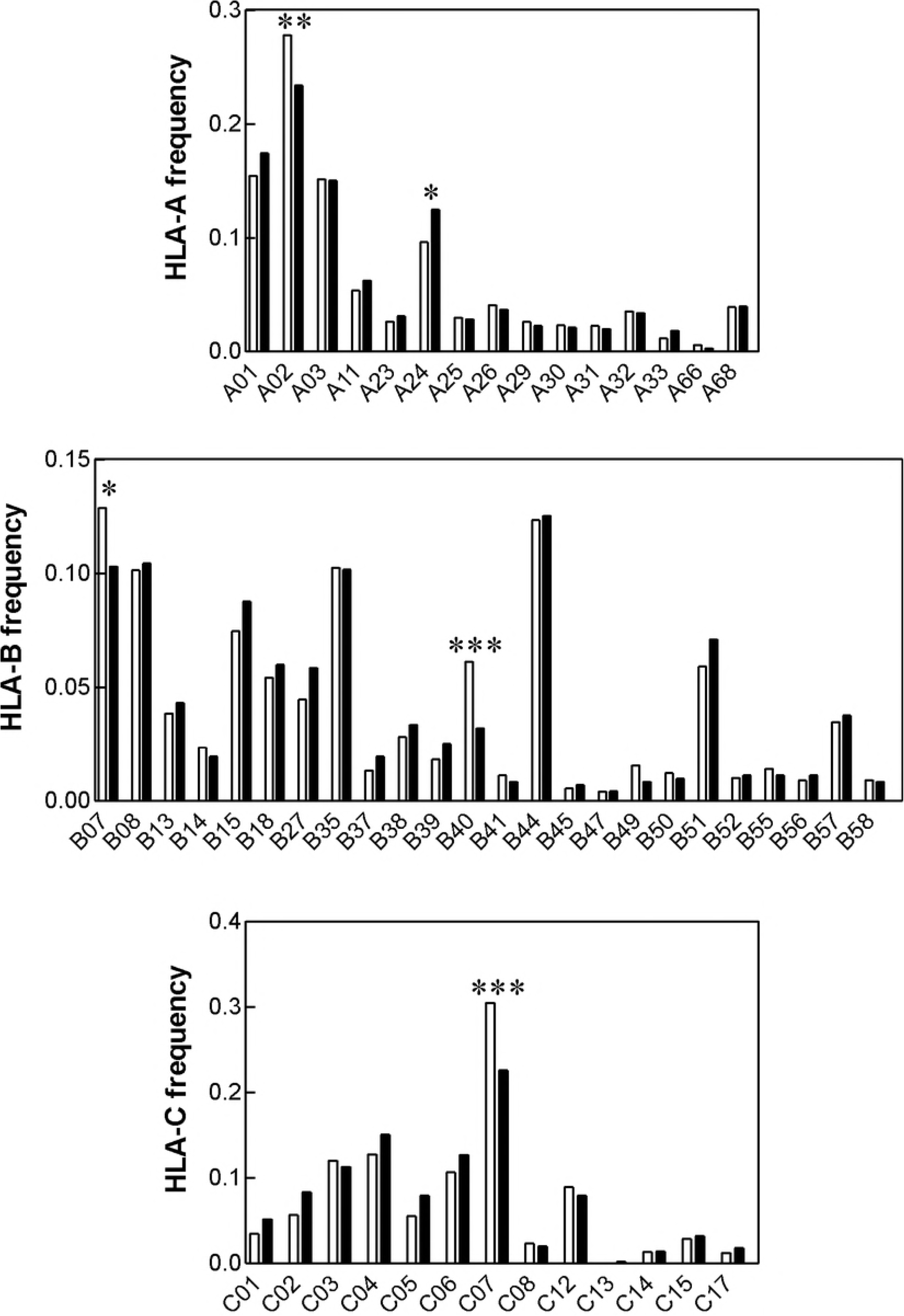
Frequency distribution of HLA class I alleles in NPMc+ AML (black bars) and in the normal population (white bars). The counts for each HLA class I allele were evaluated using contingency tables (Fischer’s exact two-tail test). Statistically significant differences between the two groups are marked with asterisks (* p < 0.05, ** p < 0.01, *** p < 0.001).

We further searched for possible impact of the HLA type on patient survival using Cox proportional hazard regression with the following risk factors: age, the presence of internal tandem duplication in *FLT3* gene (Flt3-ITD), expression of HLA-A’02, HLA-B’07, HLA-B’40 and HLA-C’07. As expected, Flt3-ITD and age were the most critical factors (with p<;0.0001 and p=0.0003, respectively). The influence of the individual HLA allelic groups did not reach statistical significance. The data are available in Supplementary Information.

We then analyzed in detail the overall survival curves for patients with and without expression of each individual HLA class I allelic group. These groups were further subdivided according to the presence or absence of Flt3-ITD which is a strong negative prognostic factor for AML. Although the curves were similar in the majority of cases (see an example for HLA-A’02 in Fig 2A), we found some differences. First, patients with B’07 expression had better survival than non-B’07 patients (Fig 2B, left). The difference was detected only for patients without Flt3-ITD (Fig 2B, right). The expression of B’40 was associated with worse survival (Fig 2C, left), in particular in Flt3-ITD-negative patients (Fig 2C, right), but the difference was not statistically significant.

**Figure 2:**
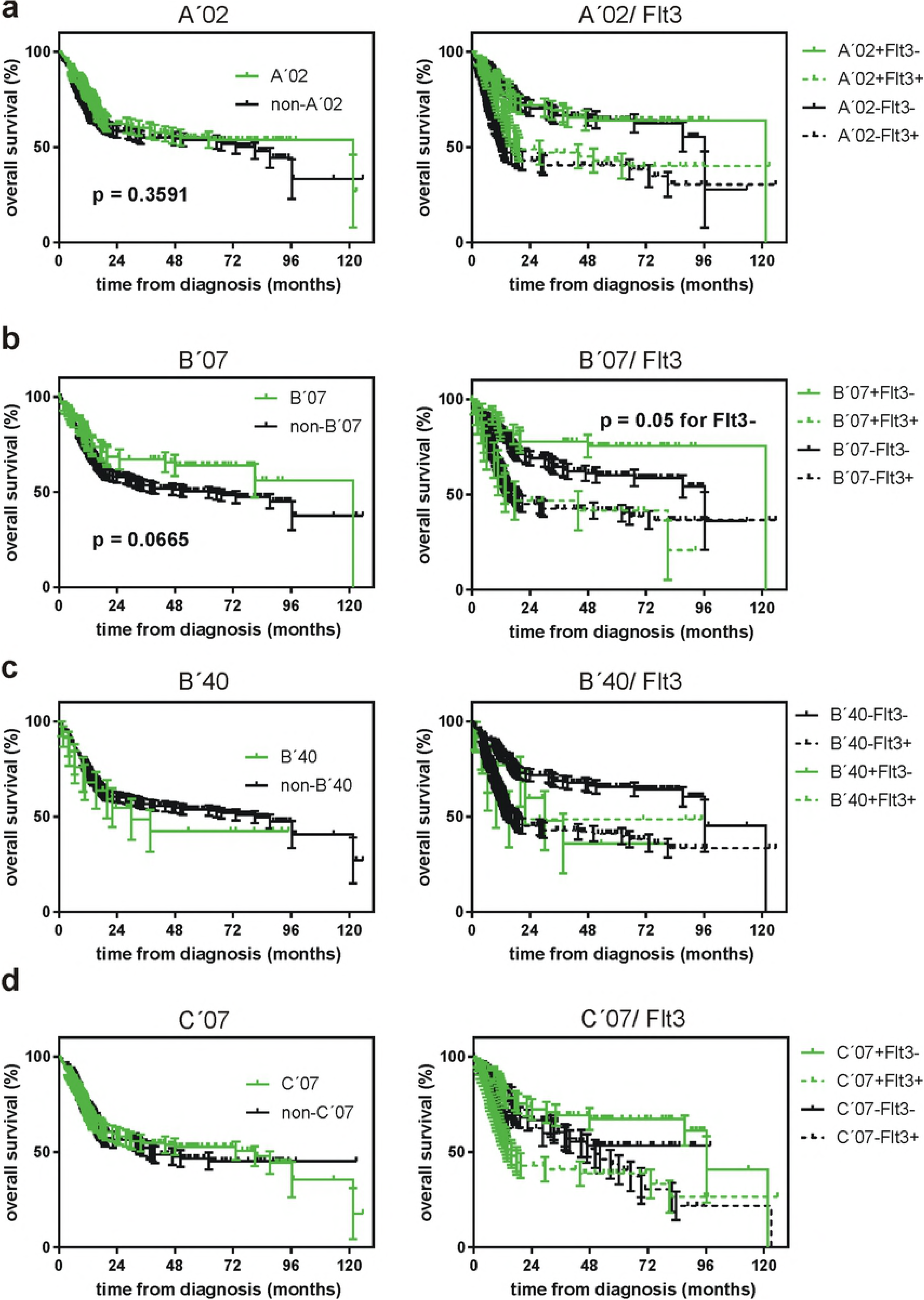
Impact of selected HLA allelic groups on the overall survival of NPMc+ AML patients. The merged cohort (N = 398) was divided according to the presence or absence of the given allelic group in the patient HLA type (left column). These groups were further subdivided according to the presence of Flt3-ITD (right column).

A complex picture was obtained for C’07 which seemed to be a positive prognostic factor in Flt3-ITD-negative group and a negative one in Flt3-ITD+ group (Fig 2D, right). The allelic group C’07 includes a number of individual alleles and the two predominant ones, C*07:01 and C*07:02, have comparable incidence. The majority of HLA data in the merged cohort were obtained by serology, which is not able to discriminate among individual alleles of a group. However, in one subcohort (Prague center, N = 94), all typings have been performed by molecular methods and we thus could analyze separately the influence of C*07:01 and C*07:02 in this subcohort (Fig 3). C*07:01, but not C*07:02 expression was associated with better survival trend although the differences were not statistically significant, probably due to the smaller number of patients (Fig 3B,C). The positive effect of C*07:01 expression was only seen in the Flt3-ITD-negative subgroup (Fig 3B, right).

**Figure 3:**
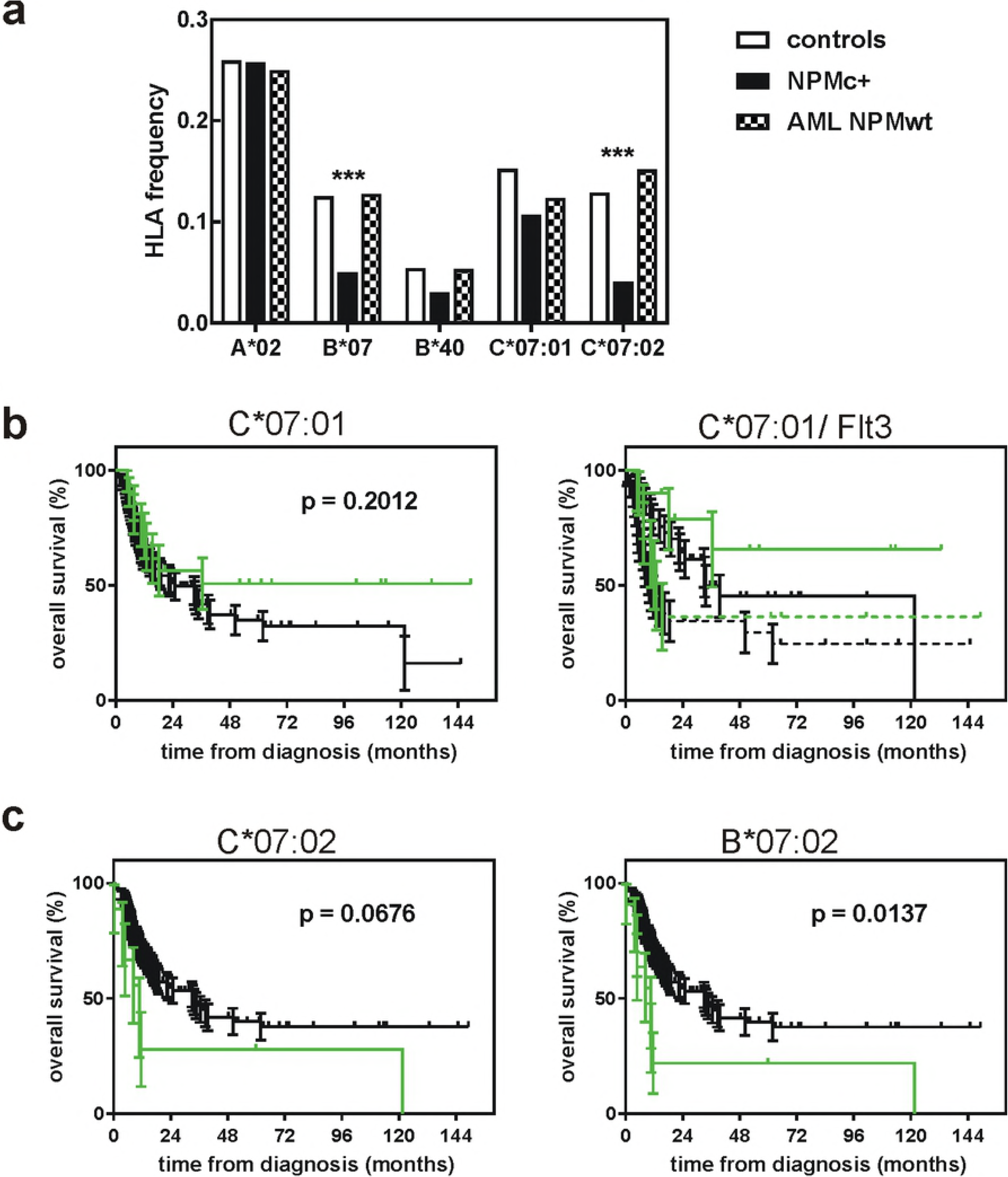
Analysis of the Prague subcohort (HLA typing by molecular genetics). A: Frequency of selected HLA class I allelic groups/alleles in a normal Czech population (white) and in AML patients with NPM1 mutation (black) or with wild-type NPM1 (checked). B: Survival curves for NPMc+ AML patients with (green) or without (black) expression of HLA-C*07:01 (left). The groups were further subdivided according to the presence or the absence of Flt3-ITD (right). C: Survival curves for NPMc+ AML patients with (green) or without (black) expression of HLA-C*07:02 (left) or HLA-B*07 (right).

Fig 3A shows the observed frequencies of selected HLA alleles in the Prague subcohort of NPMc+ AML patients together with normal values for Czech population and with values obtained from a control cohort of AML patients without mutation in *NPM1* (N = 94, labeled as AML NPMwt). B*07 allelic group (which consists mainly of one highly predominant allele, B*07:02) was underrepresented markedly in this subcohort and we detected similar decrease in C*07:02, but not in C*07:01. This is probably caused by linkage disequilibrium between these HLA genes. In fact, one of the most common ancestral haplotypes links B*07:02 (the predominant allele in B’07 allelic group) to C*07:02 [11]. In our cohort, the majority of B*07:02 expressing patients expressed also C*07:02 (19 of 20, 95%). On the other hand, in B*07-negative and C*07-positive patients, the dominant allele was C*07:01 (30 patients with C*07:01, 4 patients with C*07:02). Thus, the decrease in C*07:02 was related to that in B*07. Also, the effect of B*07 expression on the overall survival in the Prague subcohort is close to that of C*07:02 (Fig 3C) and differs from B*07 effect in the merged cohort (cf Fig 3C vs Fig 2B). On the other hand, changes associated with C*07:01 are independent of B*07. The main results of the statistical analyses are summarized in Table 2.

**Table 2.** Summary of results from the merged cohort and from the Prague subcohort

| Cohort | HLA class I frequency distribution | Effect of HLA type on the survival |
| --- | --- | --- |
| Merged<br>(N = 398) | **decrease in *HLA-A\*02* (p = 0.009)<br>*decrease in *HLA-B\*07* (p = 0.0407)<br>***decrease in *HLA-B\*40* (p = 0.0007)<br>***decrease in *HLA-C\*07* (p = 0.0001) | better survival in *HLA-B\*07+* group, specifically in the absence of *Flt3*-ITD (p = 0.05)<br>worse survival in *HLA-B\*40+* group, especially in the absence of *Flt3*-ITD (p = 0.1163) |
| Prague<br>(N = 94) | ***decrease in *HLA-B\*07* (p = 0.0007)<br>decrease in *HLA-B\*40* (p = 0.1536)<br>decrease in *HLA-C\*07:01* (p = 0.0869)<br>***decrease in *HLA-C\*07:02* (p = 0.00005) | *worse survival in *HLA-B\*07+* group (p = 0.0137)<br>better survival in *HLA-C\*07:01* group (p = 0.2012), specifically in the absence of *Flt3*-ITD |

C-terminal mutations markedly affect the intracellular localization of nucleophosmin and the majority of the mutated protein occurs in the cytoplasm whereas the wild-type form resides mainly in the nucleoli [12,13]. This could increase the exposition of NPM-derived peptides on the cell surface and trigger an immune response mediated by cytotoxic T-cells. The search for possible nucleophosmin-derived immunopeptides was performed using different prediction tools: Immune Epitope Database (IEDB) website is the most complex and provides the total score (the overall probability for each peptide to be produced and exposed in the given MHC complex) and the immunogenicity score. The binding affinity for individual HLA alleles can also be predicted using Syfpeithi or Bimas databases. The immunogenicity of the peptides was also assessed from the antigenic plot obtained using http://imed.med.ucm.es/Tools/antigenic.pl. We used the mutated NPM sequence type A as the input to find peptides with predicted high affinity to A*02, B*07, B*40 and C*07. Table 3 lists all peptides with positive IEDB total score (which have only been found for B*07 and B*40) and the best peptides that could be found for A*02. The best scores were consistently obtained for FEITPPVVL binding to B*40:01/40:02, in agreement with the marked reduction (by 50%) of B*40 incidence in NPMc+ AML patients. Promising candidate peptides (SPIKVTLATL, SPLRPQNYL) were also identified in B*07 context, with high total score from IEDB and intermediate scores from both Bimas and Syfpeithi. The predicted immunogenicity of these peptides is lower, but they both fall into antigenic determinants of NPM sequence. On the other hand, the results obtained for A*02 were controversial: several immunogenic peptides with high score were suggested from Syfpeithi search, but all of them had a negative IEDB total score and a low dissociation half time predicted by Bimas. As for C*07:01, no prediction could be obtained from Syfpeithi or from Bimas and all NPM-derived peptides had negative IEDB total scores.

**Table 3.** Predicted NPM-derived immunopeptides

| Allele | start | end | sequence | length | total score IEDB | Immunogenicity | antigenic region | Syfpeithi score | Bimas score |
| --- | --- | --- | --- | --- | --- | --- | --- | --- | --- |
| B*07:02 | 70 | 79 | SPIKVTLATL | 10 | 0.65 | -0.05054 | + | 22 | 80 |
| B*07:02 | 219 | 228 | TPRSKGQESF | 10 | 0.31 | -0.44787 | - | 18 | 4 |
| B*07:02 | 10 | 18 | SPLRPQNYL | 9 | 0.28 | -0.07898 | + | 23 | 120 |
| B*07:02 | 10 | 19 | SPLRPQNYLF | 10 | 0.12 | -0.09081 | + | 16 | - |
| B*07:02 | 13 | 23 | RPQNYLFGCEL | 11 | 0.09 | 0.10039 | + | - | - |
| B*40:01 | 92 | 100 | FEITPPVVL | 9 | 1.22 | 0.11998 | ++ | 30 | 640 |
| B*40:01 | 67 | 76 | YEGSPIKVTL | 10 | 0.92 | -0.18215 | + | 27 | 20 |
| B*40:02 | 92 | 100 | FEITPPVVL | 9 | 0.81 | 0.11998 | ++ | - | 640 |
| B*40:02 | 67 | 76 | YEGSPIKVTL | 10 | 0.59 | -0.18215 | + | - | 20 |
| B*40:01 | 91 | 100 | GFEITPPVVL | 10 | 0.45 | 0.24194 | ++ | 19 | - |
| B*40:02 | 91 | 100 | GFEITPPVVL | 10 | 0.02 | 0.24194 | ++ | - | - |
| A*02:01 | 41 | 49 | QLSLRTVSL | 9 | -1.06 | -0.03974 | + | 26 | 21 |
| A*02:01 | 288 | 296 | CLAVEEVSL | 9 | -0.79 | 0.18417 | ++ | 26 | 21 |
| A*02:01 | 75 | 83 | TLATLKMSV | 9 | -0.54 | -0.4069 | + | 25 | 70 |
| A*02:01 | 286 | 294 | DLCLAVEEV | 9 | - | 0.1913 | ++ | 25 | 6 |
| A*02:01 | 283 | 291 | AIQDLCLAV | 9 | -0.56 | -0.06333 | ++ | 24 | 39 |
| A*02:01 | 94 | 102 | ITPPVVLRL | 9 | -1.46 | 0.08518 | ++ | 23 | 1 |
| A*02:01 | 17 | 25 | YLFGCELKA | 9 | -0.37 | -0.02151 | + | 22 | 85 |
| A*02:01 | 71 | 79 | PIKVTLATL | 9 | - | 0.0546 | + | 22 | - |
| A*02:01 | 143 | 151 | SAPGGGSKV | 9 | - | -0.17022 | - | 22 | 1 |
| A*02:01 | 101 | 109 | RLKCGSGPV | 9 | - | -0.22486 | + | 21 | - |
| A*02:01 | 110 | 119 | HISGQHLVAV | 10 | - | -0.05364 | + | 27 | 0.75 |
| A*02:01 | 48 | 57 | SLGAGAKDEL | 10 | - | -0.00431 | - | 24 | 10 |
| A*02:01 | 80 | 89 | KMSVQPTVSL | 10 | -0.64 | -0.16104 | + | 23 | 54 |
| A*02:01 | 70 | 79 | SPIKVTLATL | 10 | - | -0.05054 | + | 22 | - |
| A*02:01 | 78 | 87 | TLKMSVQPTV | 10 | -0.09 | -0.46332 | + | 22 | 2.4 |
| A*02:01 | 93 | 102 | EITPPVVLRL | 10 | - | 0.09058 | ++ | 21 | - |

The best NPM-derived candidate peptides for binding to HLA-B*07, B*40 and A*02 predicted using different tools. The IEDB total score reflects the affinity of the peptide to MHC molecules, proteasome processing and TAP transport. The immunogenicity of the peptide is assessed using IEDB immunogenicity score or from antigenic plot. Syfpeithi database scores upon peptide affinity to MHC alleles. Similarly, Bimas gives the estimated half-time of dissociation for the peptide-MHC complex. No good candidate peptides were found for HLA-C*07:01.

## Discussion

Characteristic mutations in *NPM1* occur in a third of adult AML patients and they are considered to be a positive prognostic factor, at least in the absence of karyotype aberration and of other recurrent gene mutations, according to European Leukemia Net guidelines. The reason for better prognosis of NPMc+ patients is unknown and it has been suggested that anti-NPM immune response could contribute to more favorable clinical outcome of these patients [8,9]. The activity of antigen-specific cytotoxic T-lymphocytes depends on HLA type and we thus hypothesized that the individuals with suitable HLA type might be less prone to the development of NPMc+ AML. We have indeed previously identified several allelic groups of HLA class I with decreased incidence in NPMc+ AML patients, compared to the distribution in the normal population [10]. To validate our previous findings, we analyzed a larger cohort of AML patients with nucleophosmin mutation (N = 398). The decrease of B’07, B’40 and C’07 frequency was reproduced and was statistically significant, confirming a role for HLA class I type in the disease evolution (Fig 1, Table 2). In comparison with the pilot study (N = 63), we did not found differences in the frequency of HLA-B’18 and C’03 in the validation cohort. In addition, the analysis of the larger cohort indicated a decrease in HLA-A’02 frequency, which was not detected in the first cohort (Fig 1). HLA-A’02 is the preferred allelic group for the design of antigen-specific immunotherapeutical protocols as it is expressed in about a half of the Caucassian population. However, we did not observe HLA-A’02 depletion in the Prague subcohort (Fig 3A) and patients with HLA-A’02 had no survival advantage in the merged cohort (Fig 2A), as well as in the subcohorts from the individual centers. No suitable NPM-derived peptide binding to A’02 was found using theoretical predictions involving IEDB total processing score. Therefore, we believe that other HLA context should be prioritized in the search for potential target sequences although NPM-derived peptides with affinity to HLA-A’02 may also stimulate an immune response. We found the most consistent depletion in HLA-B’40 (Fig 1, Fig 3A) and, accordingly, promising NPM-derived peptides for this allelic group were predicted (Table 3). However, HLA-B’40 is not very frequent and the positive effect of the spontaneous immune response seems to be limited to the pre-diagnostic phase of the disease as the overall survival of B’40-positive patients was not better (Fig 2C).

The depletion of B’07 at diagnosis is less marked in the merged cohort (Fig 1), but a positive effect of HLA-B’07 expression was also noted at later phases (Fig 2B) and candidate NPM-derived peptides could also be found for this allelic group (Table 3). In the Prague subcohort, B’07 was much less frequent at diagnosis (Fig 3A), but the subsequent course of the disease was not better in B’07- positive group (Fig 3C). This situation is similar to that found for B’40 in the whole merged cohort (Table 2). To explain these findings, we propose the following model: an efficient immune response in the first (prediagnostic) phase of the disease, which is reflected by markedly decreased incidence of the given HLA allelic group at diagnosis, selects for cases where the immune response is inhibited or counteracted by other mechanisms. Consequently, the outcome of the patients, who have developped leukemia in spite of the predisposition for anti-NPM immune response, might be less favorable, at least in the first period of treatment. A plateau phase is often reached for longer follow-up indicating that the immune response might still help the long-term survival.

Interesting results were also obtained in the case of HLA-C’07 allelic group, which is as frequent as HLA-A’02 and has thus the potential to be used for generation of antigen-specific immune response. However, the interpretation is more complicated in this case, as this allelic group involves two alleles with comparable incidence which seem to differ in their impact. The analysis of the Prague subcohort indicates that C*07:01 is a positive prognostic factor for the overall survival (Fig 3B) in the absence of Flt3-ITD. Decreased incidence of C’07 at diagnosis is probably due more to the linkage of C*07:02 with B*07:02 than to an immune response against antigens bound to C’07. Nevertheless, in the merged cohort the decrease in C’07 is larger than that of B’07 (Fig 1) and the frequency of C*07:01 is thus probably also lowered in NPMc+ AML.

Although the prediction tools may not always identify all relevant antigens, the lack of suitable NPM-derived peptides for certain alleles may indicate the fact that the immune response is not directed against NPM, but rather against another protein which is affected by altered localization and/or signaling of the mutated NPM. Similar scenario was described for the fusion kinase BCR-ABL, which is probably not immunogenic in itself but its expression increases the immunogenicity of its downstream targets [14,15].

In general, the influence of HLA type on the overall survival is more marked in patients without Flt3-ITD (Figs 2 and 3). This correlates with the fact that patients with *NPM1* mutations have more favorable prognosis only in the absence of concomitant recurrent mutations, and especially in the absence of Flt3-ITD [16]. The negative impact of Flt3-ITD is thus usually stronger than the benefit of the presumed immune response.

## Conclusions

We have identified three distinct HLA types of NPMc+ AML patients which differ in the disease incidence and evolution (Table 2): (i) in patients expressing HLA-B’40, the presumed positive impact of the immune response occurrs mainly in the early phase of the disease and may prevent leukemia development. (ii) In patients with B*07:02/C*07:02 haplotype, the immune response would act both in the pre-diagnostic phase and during the treatment and could contribute to durable cure. (iii) The positive effect of C*07:01 expression is predominantly observed during the treatment and contributes to better overall survival. Candidate immunopeptides derived from nucleophosmin have been found for the first two groups (Table 3). In the case of C*07:01, the immune response might be directed against a different target protein. Because of their potential relevance, these findings need to be replicated in independent cohorts of patients, optimally in patients who received homogeneous AML treatment. Altogether, our analysis suggest that an immune response against nucleophosmin or against its interacting partners could explain better prognosis of NPMc+ patients and provide a rationale for attempts to explore the importance of various immunosuppressive mechanisms in this AML subgroup.

## Methods

### Pacient cohorts

All available data from patients with AML, with mutations of the type A/D/B in the gene for nucleophosmin (*NPM1*) and with known HLA class I typing results were included (total N = 398). The anonymized data were contributed from the following centers: University Hospital Carl Gustav Carus, Dresden, Germany (N = 220), Institute of Hematology and Blood Transfusion, Prague, Czech Republic (N = 94), University Hospital Brno, Czech Republic (N = 42), University of Tübingen, Tübingen, Germany (N = 25), Medical University of Silesia, Katowice, Poland (N = 17). The presence and the type of NPM1 mutation, the presence of Flt3-ITD and the HLA class I type were determined as a part of routine clinical analyses at each contributing center. In the majority of cases, HLA typing was performed using serological methods. The Prague cohort was typed by molecular genetic methods. The basic parameters of the merged cohort and of the relevant subgroups are given in Table 1.

### Ethics statement

The study has been approved by the Ethics Committee of the Institute of Hematology and Blood Transfusion of the Czech Republic as a part of the research project 16-30268A. The data were analyzed anonymously.

### Statistical analyses

The frequencies of the individual class I allelic groups in patients with mutated *NPM1* and with wild-type *NPM1* were compared with normal frequencies of these allelic groups using contingency tables (Fisher’s exact two-tailed t-test). A p-value of less than 0.05 was considered indicative of a statistically significant difference between groups. The normal frequencies were obtained from www.allelefrequencies.net. The reference cohorts used included 11407 individuals from Germany and 5099 individuals from the Czech Republic. The normal HLA class I distribution in these two countries is similar and a proportional average between German and Czech values was taken as the reference for the merged cohort. Survival curves were created and statistically evaluated using GraphPad Prism software, the Mantel-Cox test was used to assess differences between patient groups. MATLAB R2017a and SPSS were used to compute other statistics.

### Immunopeptide predictions

The MHC class I binding predictions were made on 29/9/2017 using the IEDB analysis resource Consensus tool [17] which combines predictions from ANN aka NetMHC [18,19], SMM [20] and Comblib [21]. Proteasomal cleavage/TAP transport/MHC class I combined predictor combines predictors of proteasomal processing, TAP transport, and MHC binding to produce an overall score for each peptide’s intrinsic potential of being a T-cell epitope. T-cell class I pMHC immunogenicity predictor uses amino acid properties as well as their position within the peptide to predict the immunogenicity of a class I peptide MHC (pMHC) complex. Peptides with high affinity to the individual MHC class I molecules were also searched using complementary tools: Syfpeithi database of MHC ligands (http://www.syfpheithi.de), and Bimas which ranks 8- to 10-mer peptides following a predicted half-time of dissociation from MHC molecules [22]. The immunogenicity of the peptides was also checked from the antigenic plot generated using http://imed.med.ucm.es/Tools/antigenic.pl.

## Supplementary Information

S1 Dataset for multivariate analysis

## References

[1] Teague R, Kline J. Immune evasion in acute myeloid leukemia: current concepts and future directions. Journal for ImmunoTherapy of Cancer 2013;1(1):13.

[2] Austin R, Smyth MJ, Lane SW. Harnessing the immune system in acute myeloid leukaemia. Crit Rev Oncol Hematol 2016 Jul;103:62–77.

[3] Greiner J, Hofmann S, Schmitt M, Gotz M, Wiesneth M, Schrezenmeier H, et al Acute myeloid leukemia with mutated nucleophosmin 1: an immunogenic acute myeloid leukemia subtype and potential candidate for immune checkpoint inhibition. Haematologica 2017 Dec;102(12):e499–e501.

[4] Krupka C, Kufer P, Kischel R, Zugmaier G, Lichtenegger FS, Kohnke T, et al Blockade of the PD- 1/PD-L1 axis augments lysis of AML cells by the CD33/CD3 BiTE antibody construct AMG 330: reversing a T-cell-induced immune escape mechanism. Leukemia 2016 Feb;30(2):484–491.

[5] Poh SL, Linn YC. Immune checkpoint inhibitors enhance cytotoxicity of cytokine-induced killer cells against human myeloid leukaemic blasts. Cancer Immunol Immunother 2016 May;65(5):525–536.

[6] Anguille S, Van Tendeloo VF, Berneman ZN. Leukemia-associated antigens and their relevance to the immunotherapy of acute myeloid leukemia. Leukemia 2012 Oct;26(10):2186–2196.

[7] Dvorakova D, Racil Z, Jeziskova I, Palasek I, Protivankova M, Lengerova M, et al Monitoring of minimal residual disease in acute myeloid leukemia with frequent and rare patient-specific NPM1 mutations. Am J Hematol 2010 Dec;85(12):926–929.

[8] Liso A, Colau D, Benmaamar R, De Groot A, Martin W, Benedetti R, et al Nucleophosmin leukaemic mutants contain C-terminus peptides that bind HLA class I molecules. Leukemia 2008 Feb;22(2):424–426.

[9] Greiner J, Schneider V, Schmitt M, Gotz M, Dohner K, Wiesneth M, et al Immune responses against the mutated region of cytoplasmatic NPM1 might contribute to the favorable clinical outcome of AML patients with NPM1 mutations (NPM1mut). Blood 2013 Aug 8;122(6):1087–1088.

[10] Kuzelova K, Brodska B, Fuchs O, Dobrovolna M, Soukup P, Cetkovsky P. Altered HLA Class I Profile Associated with Type A/D Nucleophosmin Mutation Points to Possible Anti-Nucleophosmin Immune Response in Acute Myeloid Leukemia. PLoS One 2015 May 20;10(5):e0127637.

[11] Dorak MT, Shao W, Machulla HK, Lobashevsky ES, Tang J, Park MH, et al Conserved extended haplotypes of the major histocompatibility complex: further characterization. Genes Immun 2006 Sep;7(6):450–467.

[12] Falini B, Mecucci C, Tiacci E, Alcalay M, Rosati R, Pasqualucci L, et al Cytoplasmic nucleophosmin in acute myelogenous leukemia with a normal karyotype. N Engl J Med 2005 Jan 20;352(3):254–266.

[13] Brodska B, Kracmarova M, Holoubek A, Kuzelova K. Localization of AML-related nucleophosmin mutant depends on its subtype and is highly affected by its interaction with wild-type NPM. PLoS One 2017 Apr 6;12(4):e0175175.

[14] Brauer KM, Werth D, von Schwarzenberg K, Bringmann A, Kanz L, Grunebach F, et al BCR-ABL activity is critical for the immunogenicity of chronic myelogenous leukemia cells. Cancer Res 2007 Jun 1;67(11):5489–5497.

[15] Scheich F, Duyster J, Peschel C, Bernhard H. The immunogenicity of Bcr-Abl expressing dendritic cells is dependent on the Bcr-Abl kinase activity and dominated by Bcr-Abl regulated antigens. Blood 2007 Oct 1;110(7):2556–2560.

[16] Thiede C, Koch S, Creutzig E, Steudel C, Illmer T, Schaich M, et al Prevalence and prognostic impact of NPM1 mutations in 1485 adult patients with acute myeloid leukemia (AML). Blood 2006 May 15;107(10):4011–4020.

[17] Kim Y, Ponomarenko J, Zhu Z, Tamang D, Wang P, Greenbaum J, et al Immune epitope database analysis resource. Nucleic Acids Res 2012 Jul;40(Web Server issue):W525–30.

[18] Nielsen M, Lundegaard C, Worning P, Lauemoller SL, Lamberth K, Buus S, et al Reliable prediction of T-cell epitopes using neural networks with novel sequence representations. Protein Sci 2003 May;12(5):1007–1017.

[19] Lundegaard C, Lamberth K, Harndahl M, Buus S, Lund O, Nielsen M. NetMHC-3.0: accurate web accessible predictions of human, mouse and monkey MHC class I affinities for peptides of length 8-Nucleic Acids Res 2008 Jul 1;36(Web Server issue):W509–12.

[20] Peters B, Sette A. Generating quantitative models describing the sequence specificity of biological processes with the stabilized matrix method. BMC Bioinformatics 2005 May 31;6:132.

[21] Sidney J, Assarsson E, Moore C, Ngo S, Pinilla C, Sette A, et al Quantitative peptide binding motifs for 19 human and mouse MHC class I molecules derived using positional scanning combinatorial peptide libraries. Immunome Res 2008 Jan 25;4:2-7580-4-2.

[22] Parker KC, Bednarek MA, Coligan JE. Scheme for ranking potential HLA-A2 binding peptides based on independent binding of individual peptide side-chains. J Immunol 1994 Jan 1;152(1):163– 175.

